# Electrical Coupling between Parvalbumin Basket Cells is Reduced after Experimental Status Epilepticus

**DOI:** 10.1101/2023.09.27.559804

**Authors:** Jiandong Yu, Vijayalakshmi Santhakumar

**Affiliations:** Department of Neurosurgery, the First Affiliated Hospital of Bengbu Medical College, Bengbu, China; Department of Pharmacology, Physiology and Neuroscience, Rutgers New Jersey Medical School, Newark, New Jersey 07103; Department of Molecular, Cell and Systems Biology, University of California Riverside, Riverside, California 92521

**Keywords:** electrical synapses, parvalbumin, connexin 36, acquired epilepsy, interneuron, synchrony, inhibition

## Abstract

Acquired epilepsies, characterized by abnormal increase in hypersynchronous network activity, can be precipitated by various factors including brain injuries which cause neuronal loss and increases in network excitability. Electrical coupling between neurons, mediated by gap junctions, has been shown to enhance synchronous neuronal activity and promote excitotoxic neurodegeneration. Consequently, neuronal gap junctional coupling has been proposed to contribute to development of epilepsy. Parvalbumin expressing interneurons (PV-INs), noted for their roles in powerful perisomatic inhibition and network oscillations, have gap junctions formed exclusively by connexin 36 subunits which show changes in expression following seizures, and in human and experimental epilepsy. However, only a fraction of the connexin hemichannels form functional connections, leaving open the critical question of whether functional gap junctional coupling between neurons is altered during development of epilepsy. Using a pilocarpine induced status epilepticus (SE) model of acquired temporal lobe epilepsy in rat, this study examined changes in electrical coupling between PV-INs in the hippocampal dentate gyrus one week after SE. Contrary to expectations, SE selectively reduced the probability of electrical coupling between PV-INs without altering coupling coefficient. Both coupling frequency and coupling coefficient between non-parvalbumin interneurons remained unchanged after SE. The early and selective decrease in functional electrical coupling between dentate PV-INs after SE may represent a compensatory mechanism to limit excitotoxic damage of fast-spiking interneurons and network synchrony during epileptogenesis.

## Introduction

Acquired temporal lobe epilepsy (TLE) can develop as a consequence a variety of brain insults and is characterized by abnormal increases in hippocampal network excitability and synchrony. The process of epileptogenesis involves cellular and circuit alterations in the hippocampal dentate gyrus (DG) including neuronal loss, altered neuronal physiology and synaptic connectivity. Apart from synaptic connections, gap junctional connections between neurons, mediated by connexin hemichannles provide an alternative unique mechanism for direct rapid electrical and small molecule communication between cells (Alcami and Pereda, 2019). Gap junctions have been observed both between neurons and between glia with cell-type specific difference is the underlying connexin subtypes. Interestingly, heterotypic neuro-glial coupling has also been observed under certain conditions. Among neurons, gap junctional coupling has been established, and most extensively examined, in inhibitory neurons including parvalbumin expressing interneurons (PV-INs) in which electrical coupling is mediated by connexin 36 (Cx36) hemichannels (Hormuzdi et al., 2001). The electrical coupling mediated by gap junctions can modify neuronal resistive and capacitive properties and support transmission of sub-threshold currents including synaptic currents and action potential spikelets (Alcami and Pereda, 2019). Consequently, gap junctions can dynamically regulate synaptic integration, excitability and synchrony of coupled neurons in circuits. Consistently, electrical coupling between fast-spiking PV-INs in the DG has been shown to support network synchrony (Bartos et al., 2002). Due to its potential to promote synchrony, gap junctions have been proposed to contribute to seizures and epilepsy (Mylvaganam et al., 2014). Supporting this hypothesis, blocking or genetically deleting Cx36 has been shown to reduce acute chemically evoked seizure-like events in hippocampal circuits (Zsiros et al., 2007). However, whether functional electrical coupling between interneuron subtypes is enhanced in epilepsy has not been examined.

Apart from its electrophysiological function in promoting synchrony, the ability of gap junctions to facilitate exchange of small molecules can promote excitotoxic neuronal injury (Wang et al., 2012) which could also contribute to epileptogenesis. While several studies have examined changes hippocampal Cx36 mRNA and protein expression, the results have been variable depending on time after seizures and experimental model (Beheshti et al., 2010; Mylvaganam et al., 2014). Regrettably, these studies lack cell type specificity and whether interneuron subtypes show differential changes in connexin expression in epilepsy has not been examined. Nevertheless, since only a fraction of connexin hemichannels form functional connections and multiple intracellular processes can regulate gap junctional coupling (Alcami and Pereda, 2019), connexin expression may not directly translate to functional electrical coupling. The relatively impermeability of Cx36 to most dyes used to assess coupling has limited the ability to assess changes in interneuronal gap junctional coupling. Here we undertook paired recordings from DG fast-spiking PV-INs and non-fast spiking accommodating interneurons one week after pilocarpine induced status epilepticus (SE) in rat (Yu et al., 2016) to directly examine whether electrical coupling between dentate interneurons is altered during epileptogenesis.

## Methods

### Experimental animals

Wistar rats pups with nursing dams were purchased from Charles River (USA) and weaned at postnatal day 21. Young adult male rats aged 25-28 days were used in experiments. Animals were group house with 2-4 rats/cage, under a 12-h/12-h light/dark cycle with controlled temperature (20–26 °C), humidity (50 ± 20 %) and *ad libitum* access to food and water under protocols approved by the Institutional Animal Care and Use Committee at Rutgers-NJMS, Newark, NJ. All experimental methods conform to the ARRIVE guidelines.

### Pilocarpine-induced seizures

Animals were pretreated scopolamine (1 mg/kg, s.c) dissolved in normal saline (1 mg/mL). This was followed after 30 minutes by injection with pilocarpine (350 mg/kg, i.p.) dissolved in normal saline (150 mg/mL). Diazepam (5 mg/mL, i.p.) was used to terminate the status epilepticus (SE) one hour after the first stage 3 or greater seizures on the Racine scale (Yu et al., 2016). Age-matched littermate controls received saline instead of pilocarpine and diazepam after 2 hours.

### Paired patch-clamp recordings

One week (7-10 days) after SE, experimental animals were euthanized under 1.5% isoflurane. Transverse hippocampal slices (250μ m) were prepared in the ice-cold sucrose artificial cerebrospinal fluid (aCSF) containing in mM, 75 mM sucrose, 85 mM NaCl, 25 D-glucose, 24 NaHCO_3_, 4 MgCl_2_, 2.5 KCl, 1.25 NaH_2_PO_4_, and 0.5 CaCl_2_ with 95%O_2_ and 5% CO_2_. Slices were incubated in recording aCSF containing, in mM, 126 NaCl, 26 NaHCO3, 10 D-glucose, 2.5 KCl, 2 CaCl_2_, 2 MgCl_2_ and 1.25 NaH_2_PO_4_ bubbled with 95%O_2_ and 5% CO_2_, first at 33±1°C for 30 min, and then at room temperature.

Dual whole cell recordings were obtained at 33±2ºC in recording aCSF. Large triangular neurons in the hilar-granule cell layer-border were patched under IR-DIC visualization using a Nikon FN-1 microscope (40X water-objective). Recording electrodes (3-4 MΩ) contained (in mM) 70 KCl, 70 K-gluconate, 10 HEPES, 2 MgCl2, 0.2 EGTA, 2 Na-ATP, 0.5 Na-GTP, 10 phosphocreatine and 0.2% biocytin. Intrinsic active and passive properties were determined from responses to 1.5 sec current injections from a holding potential of -70 mV. Neurons with non-adapting, high frequency firing, and low input resistance (<150 MΩ) were considered putative PV basket cells. Post-hoc biocytin immunostaining and morphological analysis was used to definitively identify PV-INs based on co-labeling for PV and biocytin and axons in the granule cell layer. Neurons with adapting firing without stuttering, and input resistance over 150 MΩ were considered non-PV interneurons and typically showed dendrites in the hilus and axonal projections in the molecular layer (Yu et al., 2016). In unconnected neuron pairs used to estimate connection probability, cell types were defined based on physiology. Data were collected using pClamp 10 (Molecular Devices, Palo Alto, CA), sampled at 10 kHz and filtered at 3 kHz, analyzed with Clampfit 10.4.

Recorded slices were fixed in 4% paraformaldehyde, incubated overnight with anti-parvalbumin antibody (PV-28, 1.5:1000, polyclonal rabbit) in 0.3% Triton X-100 and 2% normal goat serum. Immunoreactions were revealed using Alexa 488-conjugated secondary antibody. Biocytin was revealed using Alexa 594-conjugated streptavidin (1:1000). Sections were visualized and imaged using a Zeiss LSM510 confocal microscope (0.5 NA, 20X objective).

### Statistics

Statistical analysis was performed with Origin 8. Statistical significance was tested was two-tailed Mann-Whitney tests, and chi-test as appropriate. All data are shown as mean ± SEM. The significance level was set to P < 0.05.

## Results

### Gap junctional coupling between parvalbumin interneurons is reduced after SE

Fast-spiking parvalbumin interneurons (PV-INs) are known to have extensive gap junctional connectivity (Bartos et al., 2002). Dentate PV-INs, which include perisomatically projecting basket cells and axon-initial segment targeting axo-axonic cells, are also relatively spared from early cell loss and functional changes early after pilocarpine induced SE (Yu et al., 2016). We undertook paired recordings from parvalbumin basket cells to determine functional changes in electrical coupling among PV-INs in rats one week after pilocarpine induced SE and age-matched saline-treated controls (figure 1A). Basket cells were targeted based on location at the border of granule cell layer and hilus and definitively identified based on post hoc morphology of biocytin filled neurons exhibiting basket-like axon projections in the granule cell layer and PV expression in the somata (figure 1B). Recorded neurons were physiologically classified as fast-spiking based on non-adapting firing in response to positive current injections (figure 1C). As illustrated in figure 1C, negative and positive current injections in one fast-spiking PV-IN elicited a corresponding hyperpolarization or depolarization in the electrically connected PV-IN. Moreover, firing in one PV-IN elicited spikelets in the connected PV-IN and the connections were reciprocal. In recordings from fast-spiking PV-IN pairs in control rats, the probability of detecting electrical connections was 44% (22 of 50 pairs) and the coupling coefficient (defined as the ratio between steady state voltage change in coupled neuron and the voltage in the neuron receiving current injection) was 0.03±0.01 (figure 1D, E, n=20 pairs). These values are consistent with the recordings from other labs (Bartos et al., 2002). Surprisingly, we identified a significant reduction in the coupling probability (figure 1D, 23%, 17 of 74 pairs, P<0.01, χ^2^-test) one week after SE. However, the coupling coefficient was not different between control and post-SE rats (figure 1E, 0.03±0.01, n= 17 control; 0.03±0.01, 14 SE-early pairs P>0.05, Mann-Whitney test).

**Figure 1.**
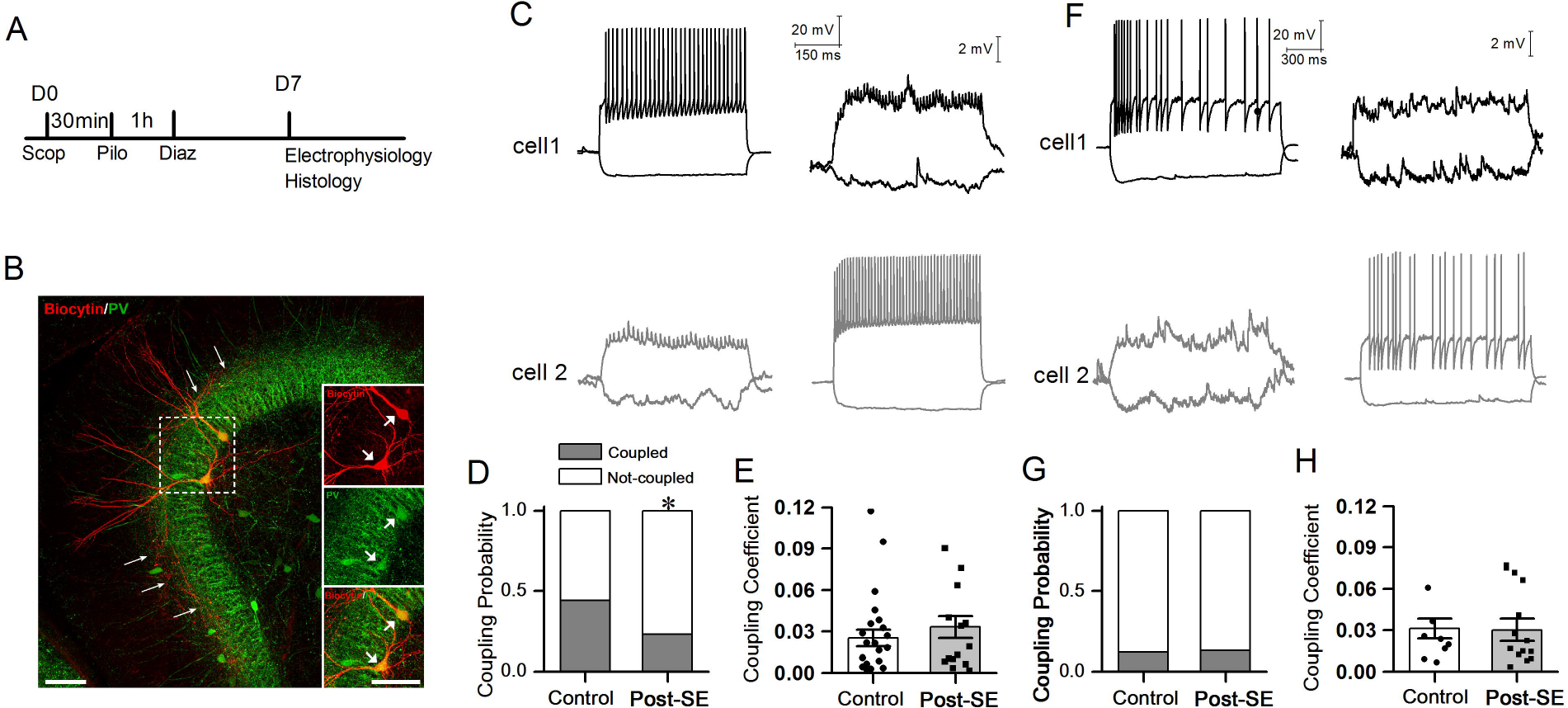
Selective reduction in functional electrical coupling between fast-spiking parvalbumin interneurons one week after SE. (A) Schematic of experimental design illustrates timeline for administration of scopolamine (scop) followed by pilocarpine (Pilo). Diazepam (Diaz) was administered at 1 hour followed by euthanasia for slice physiology after 1 week. (B) Right panel illustrates confocal images of electrically coupled dentate parvalbumin basket cells show the biocytin filled soma and dendrites are labeled with PV. Arrows denote the biocytin labeled neurons have with axons in the cell layers. Arrows in the insert images show the somatic co-localization of biocytin and PV. Scale bars represents 50 μm. (C) Example traces illustrate reciprocal electrical coupling between fast spiking parvalbumin interneurons. Note that both neurons show fast spiking firing pattern (upper right and lower left). The ability of firing in one neuron to evoke spikelets in coupled neuron is illustrated. (D-E) Summary plots of the probability of electrical coupling between fast spiking parvalbumin interneurons (D) and the coupling coefficients in these pairs (E) in control and post-SE rats. (F) Example traces illustrate reciprocal electrical coupling between non-fast spiking interneurons. (G-H) Summary plots of the coupling probability of electrical coupling (G) and coupling coefficients in the non-fast spiking interneuron pairs (H) in control and post-SE rats. * indicates p<0.05 by χ^2^-test.

### Gap junctional coupling between non-fast spiking interneurons is not altered after SE

We next examined electrically connected neuronal pairs between non-fast spiking interneurons, identified by the presence of spike frequency adaptation during positive current injection (figure 1F). In paired recordings, non-fast spiking interneurons showed considerably lower electrical coupling than PV-INs (PV-INs, 44%, 22 of 50 pairs; AC-INs, 12%, 12 of 99 pairs, P<0.01, χ^2^-test). However, the probability of detecting electrical connections between non-fast spiking interneurons in rats 1 week after SE was not different from controls (figure 1G, Control: 12%, 12 of 99 pairs, post-SE: 13.4%, 17 of 126 pairs, P>0.05, χ^2^-test) rats. Additionally, the coupling coefficient in non-fast spiking neuron pairs was not different between control and SE groups (figure 1H, control 0.025±0.01, n=8 pairs; post-SE 0.033±0.01, n=14 pairs, P>0.05, Mann-Whitney test). Together, our data showed that electrical coupling between dentate fast-spiking PV-INs is selectively reduced early after SE, while gap junctional coupling between accommodating interneurons remains unchanged.

## Discussion

This study investigated changes in functional electrical connectivity between fast-spiking PV-INs in the dentate gyrus one week after SE in an experimental model of epilepsy. Contrary to expectations that gap junctional coupling would be increased after SE, we identified a decrease in the probability of electrical coupling between PV-INs after SE. This decrease is selective for PV-INs and the coupling probability in non-fast spiking interneurons was unchanged after SE. The coupling coefficient, representing the strength of electrical coupling between either interneuron subtype was not altered after SE. To our knowledge, these results are the first to identify a functional change in gap junctional coupling between interneurons in the latent period of epileptogenesis.

Spread of activity in neuronal networks is largely mediated by chemical synapses. In addition, bidirectional electrical synapses formed by gap junctions between neurons have been shown to enhance network synchrony. Uniquely, gap junctions from intercellular channels between the cytoplasm of connected cells and allow for both ions and small molecules to move between coupled cells. Gap junctions are found between neuronal dendrites, particularly in inhibitory neurons, and between glia but have also been observed between dentate granule cell axons and at certain chemical synapses (Alcami and Pereda, 2019). Both neuronal and glial gap junction can contribute to changes in network activity with neuronal gap junctions proposed to enhance synchrony, while astrocytic gap junctions contributing to glucose delivery and supporting potassium clearance during intense neuronal activity (Mylvaganam et al., 2014). Although single-cell RT-PCR has identified multiple connexin subtypes in neurons, connexin 36 underlies electrical coupling between PV-INs (Hormuzdi et al., 2001). Since neuronal gap junctions can promote synchrony they have been proposed to contribute to epilepsy (Mylvaganam et al., 2014). Moreover, since PV-INs shape network oscillations, increased gap junctional coupling between PV-INs is ideally suited to increase network synchrony. Consequently, studies have examined hippocampal expression of Cx36, the major connexin subtype expressed in PV-INs, in experimental epilepsy have found that the changes vary based on seizure model, time after SE, brain region and cell type (reviewed in Mylvaganam et al., 2014). Although it has been recognized that changes in Cx36 expression may not translate to electrical coupling which can be transient (Alcami and Pereda, 2019; Mylvaganam et al., 2014), functional assays of intraneuronal electrical coupling have been challenging due to the inability of intraneuronal Cx36 channels to support dye coupling. Our studies overcome this limitation by adopting paired recordings from dentate interneurons in a model of experimental epilepsy and demonstrate a decrease in probability of electrical coupling between PV-INs. In contrast, non-fast spiking accommodating interneurons, which included hilar commissural-association pathway (HICAP) cells, with axons in the inner molecular layer, and total-molecular layer (TML) cells (Yu et al., 2016) showed considerably lower electrical coupling than PV-INs which remained unchanged after SE. However, coupling coefficient, representing the strength of connection was not altered in either cell type indicating that gap junctional characteristics remined unchanged. In contrast, the reduction in coupling probability likely reflects cell specific intracellular regulation of Cx36 coupling in PV-INs.

The several inhibitory neuron subtypes in the hippocampus are distinguished by their morphology, intrinsic physiology, synaptic connectivity, function as well as plasticity in disease. Interneuron subtypes also differ in the extent of electrical connectivity and the connexin subtypes that underlie the coupling (Venance et al., 2000). PV-INs have been shown to exhibit higher gap junctional interconnectivity than other interneuron subtypes in the rodent hippocampus. The connection probability and coupling coefficient in our control data is consistent with prior studies. Here we find that gap junction between PV-INs was reduced while those between non-fast spiking interneurons remains unchanged after SE. This is different from the post SE changes in synaptic connections which remain unchanged between PV-INs but are strengthened between non-fast spiking interneurons (Yu et al., 2016). It is possible that the functional uncoupling of PV-INs occurs despite an increase in Cx36 expression and could be an attempt to compensate for increase in network synchrony. Moreover, reducing gap junctional coupling would be expected to enhance neuronal excitability by modifying passive membrane properties (Alcami and Pereda, 2019), suggesting that the reduction in PV-IN electrical coupling after SE would support perisomatic inhibition. It is possible that, at physiological levels, gap junctional coupling has greater impact on neuronal passive properties than on synchrony.

The gap junction channels between neurons not only allow ions mediating electrical signaling but also small molecules including second messengers that contribute to cell death (Wang et al., 2012). Increases in network excitability after SE are known to contribute to excitotoxic neuronal loss including extensive loss of dentate hilar excitatory and inhibitory neurons. Signals associated with glutamate excitotoxicity including Ca^2+^ and IP_3_ can propagate to gap junction coupled interneurons and induce cell death (Wang et al., 2012). Here we find that gap junctional connectivity among dentate PV-INs which are less prone to loss after SE (Sun et al., 2007) is decreased. This raises the possibility that uncoupling of PV-INs after SE could limit their excitotoxic loss during epileptogenesis.

In conclusion, our paired recording data demonstrate that PV-INs and not non-fast spiking interneurons in the dentate gyrus show reduced gap junctional coupling after SE. This decrease in junctional coupling has the potential to augment PV-IN excitability while limiting excitotoxic loss of coupled neurons and could be a compensatory mechanism to maintain perisomatic inhibition and limit network synchrony.

## Abbreviations

Cx36: Connexin 36
DG: Dentate Gyrus
IN: Interneuron
PV: Parvalbumin
PV-IN: Parvalbumin Interneuron
SE: Status Epilepticus
TLE: Temporal Lobe Epilepsy

## Author contributions

V.S. and J.Y. designed experiments. J.Y. conducted the experiments. V.S. and J.Y. wrote the manuscript. V. S. Secured funding.

## Data and materials availability

Materials are available upon request.

